# Humoral correlates of protection in a mouse model of echovirus infection

**DOI:** 10.1101/2025.09.24.678383

**Authors:** Patrick S. Creisher, James W. Abell, Kaila A. Cooley, Carolyn B. Coyne

**Author notes:** To whom correspondence should be addressed: Carolyn Coyne, PhD, 3130 Medical Sciences Research Building III, 3 Genome Court, Durham, NC 27710, USA.

## Abstract

Echoviruses commonly infect humans and can cause severe outcomes, including meningitis and liver failure, especially in neonates and immunocompromised individuals. Although recent progress has been made in understanding acute pathogenesis and innate immunity to echoviruses, adaptive immune responses remain poorly defined, in part due to the lack of an appropriate small animal model of infection. Here, we developed a translational mouse model of echovirus infection using hFc mice, which express human FcRn, the echovirus receptor, while maintaining functional IgG circulation. Primary echovirus infection caused acute disease and elicited virus-specific nAbs, IgG, and IgM only in hFc mice, which express human FcRn and support IgG transport, but not in parental Tg32 mice that lack IgG transport. These humoral responses correlated with protection against a homologous lethal-dose challenge. The protective role of these antibodies was confirmed by passive transfer of immune serum, which conferred complete, sterilizing protection. Statistical modeling identified E5-specific nAb, total IgG, and IgG1 titers as the strongest predictors of survival following infection, identifying novel correlates of protection. These findings establish hFc mice as a robust translational model for dissecting echovirus adaptive immunity, define the humoral response to infection, and identify correlates of protection to guide future vaccine development and preclinical evaluation.

## 1 Introduction

Enteroviruses cause a wide spectrum of human diseases, including aseptic meningitis, paralysis, myocarditis, and acute liver failure^1–4^ and are a leading cause of hospitalization in neonates and infants^5; 6^. For example, in the United States, enteroviruses have been estimated to account for 20% of febrile hospitalizations and 25% of suspected sepsis diagnoses in infants <90 days old ^7; 8^. Beyond acute illness, enterovirus infections have been associated with long-term sequelae such as an increased risk of type 1 diabetes and dilated cardiomyopathy^9–12^. Among enteroviruses, echoviruses are particularly clinically relevant, especially among neonates. In neonatal intensive care units, echovirus infections account for 15-30% of nosocomial viral infections and can have a case fatality rate exceeding 25%^13–15^. Despite the public health relevance, echoviruses remain among the least studied enteroviruses, with no licensed vaccines or antiviral therapeutics^16–18^. For example, although monoclonal antibodies targeting echovirus 30 have been identified^19^, testing of *in vivo* efficacy has been limited without a robust mouse model.

Mouse models are critical for evaluating viral pathogenesis and immune responses. Live virus challenge, often not feasible in humans, enables testing of antiviral and vaccine efficacy and the identification of potential biomarkers and correlates of protection^20^. Unlike other enteroviruses such as coxsackievirus B (CVB) or enterovirus D68 (EVD68), echoviruses cannot productively infect mouse cells or establish infection in wild-type mice^21–23^ as echoviruses require the human, but not mouse, neonatal Fc receptor (FcRn) for entry and infection.^23; 24^ The identification of human FcRn as the pan-echovirus receptor^23^ enabled the use of Tg32-hFcRn (Tg32) mice, in which mouse FcRn has been replaced with human FcRn under the native promoter^25^, to evaluate echovirus pathogenesis. Tg32 mice are permissive to echovirus infection and recapitulate key aspects of human infection, including meningitis and hepatotropism^26; 27^. Using Tg32 mice, we previously investigated the innate immune response to echovirus *in vivo* and identified type I interferon (IFN) signaling as critical for controlling acute echovirus infection. For example, hepatocyte-specific type I IFN signaling limits liver infection and prevents liver failure as well as mortality^28^, highlighting that Tg32 mice are a valuable model to study innate immune responses to echoviruses.

Despite these advances, our understanding of the adaptive immune responses to echoviruses is limited. Clinical evidence suggests a critical role for antibodies, as patients with genetic agammaglobulinemia or those receiving B cell-depleting immunosuppressive therapy are susceptible to severe and prolonged echovirus infections^29; 30^. However, the specific mechanisms of protection, such as the role of neutralizing antibodies (nAbs) and specific isotypes, remain unknown. A key limitation of Tg32 mice is impaired antibody homeostasis. Because serum IgG maintenance depends on FcRn-mediated recycling, and human FcRn binds mouse IgG poorly, Tg32 mice have reduced serum IgG levels and impaired humoral responses^31; 32^. To overcome this limitation, Tg32 mice were further engineered to produce chimeric IgG1 with mouse Fab regions and the human Fc domain^32^. These modified mice, referred to as hFc mice, exhibit improved IgG recycling and more robust antibody responses to antigen exposure.^32^

In this study, we used the hFc mouse model to compare infection and immune responses with those of the parental Tg32 mice, demonstrating its utility for defining correlates of protection and supporting the development of echovirus vaccines and therapeutics. Echovirus infection of hFc mice elicited a robust serum nAb response that correlated with survival, establishing humoral correlates of protection for preclinical testing. We identified echovirus-specific nAbs, chimeric IgG1, and mouse IgG2 titers at the time of challenge as the most robust correlates of protection. Their causal role was confirmed by passive transfer of immune serum, which conferred sterilizing protection against disease and death in naïve mice. These findings highlight the utility of hFc mice for defining adaptive immune responses to echoviruses, identifying correlates to guide vaccine design, and providing a challenge model for preclinical vaccine evaluation.

## 2 Materials and Methods

### 2.1 Cells and viruses

HeLa cells (clone 7B; provided by Dr. Jeffrey Bergelson, Children’s Hospital of Philadelphia) were cultured in HeLa media [MEM (Corning #10-010-CM) supplemented with 5% fetal bovine serum (Gibco #A56707-01), 1× nonessential amino acids (Cytiva #SH30238.01), and 100 U/mL penicillin & 100 μg/mL streptomycin (Gibco #15140–122)] at 37 °C. Viral seed stocks of E5 (Noyce strain; ATCC #VR-35) and CVB3 (H3 strain; generated from an infectious clone provided by Dr. J Lindsay Whitton^21; 33^) were propagated on HeLa cell monolayers and used to prepare purified viral working stocks for experiments, as described previously^26; 34^. Briefly, HeLa cell monolayers were infected, and after 48-96 h, supernatant and cells were collected, freeze-thawed thrice to lyse cells, treated with Triton X-100 (final concentration of 0.5%; Sigma #X100-500ML), and clarified by ultracentrifugation in a Sorvall wX+ SW28 rotor at 7200 RPM for 20 min at 16°C. Virus-containing supernatant was then treated with SDS (final concentration of 0.75%; RPI #L22040-1000) and purified by ultracentrifugation in a Sorvall wX+ SW28 rotor at 27000 RPM for 3 h at 16°C over a 30% sucrose cushion. The virus pellet was resuspended in phosphate buffered saline (PBS; Corning #21-031-CM), aliquoted, and stored at −80 °C. Viral titers were determined by plaque assay on HeLa cells,^34^ and stock purity confirmed by metagenomic next-generation sequencing^35^. To generate inactivated virus for enzyme-linked immunosorbent assays (ELISA), E5 was propagated in the presence of neutral red (10 μg/mL; ThermoScientific #229810250) and purified as above under semi-dark conditions. Purified neutral red E5 was transferred to a 60 mm polystyrene petri dish and inactivated on a lightbox at room temperature for 45 min^36; 37^. The concentration of inactivated virus was determined using the Pierce BCA Protein Assay Kit (Thermo Fisher Scientific #23227) and inactivation was confirmed by plaque assay on HeLa cells^34^.

### 2.2 Experimental mice

Male and female B6.Cg-*Fcgrt^tm1Dcr^*Tg(FCGRT)32Dcr/DcrJ (Tg32; JAX #014565)^25^ and B6.Cg-Tg(FCGRT)32Dcr *Fcgrt^tm1Dcr^ Ighg1^em^*^2^*^(IGHG^*^1^*^)Mvw^*/MvwJ (hFc; JAX #029686)^32^ were purchased from the Jackson Laboratory and either used directly for experiments or maintained as breeders to produce mice for experiments. Mice were housed under standard biosafety level 2 housing conditions (2-5 mice per cage) with *ad libitum* food and water and were acclimated to water bottles for at least 96 h prior to experiments. Monitoring and procedures for animal experiments were performed consistently at the same time of day. All animal experiments were approved by the Duke University Animal Care and Use Committee (protocol A012-24-01) and conducted in accordance with the relevant guidelines and regulations.

### 2.3 Sublethal-dose infection, serum collection, and lethal-dose challenge

Male and female mice (6-10 weeks of age) were intraperitoneally (IP) infected with 10^3^ PFU of E5 in 50 μL of PBS or mock-inoculated with 50 μL of PBS. Mice were monitored daily, and body mass recorded for 14 days after infection or until tissue collection. Mice that were moribund or had lost >25% of body mass since the day of infection were euthanized by isoflurane inhalation overdose (drop method) followed by cardiac exsanguination. At days 14, 21, 28, 35, and 42 post-infection, blood was collected from the submandibular vein under isoflurane anesthesia. Serum was separated by centrifugation (9600 × g for 30 min at 4°C) and stored at −80°C until analysis. At day 42 post-infection, surviving mice were challenged with 10^6^ PFU of E5 or CVB3 in 50 μl of PBS via IP injection, and monitored for 14 days for morbidity (as measured by body mass change) or mortality, with humane endpoints as described above.

### 2.4 Tissue collection and homogenization

A subset of infected or mock-inoculated mice were euthanized by isoflurane inhalation overdose (drop method) followed by cardiac exsanguination at 3 days post sublethal-dose infection or 3 days post lethal-dose challenge. Whole blood, liver, and pancreas were collected and flash frozen on dry ice. Flash-frozen pancreases and liver median lobes were weighed and homogenized in 1:10 weight per volume of PBS using 2.8 mm ceramic bead-beating tubes (Revvity #19-628) in an Omni International Bead Ruptor Elite instrument (4m/s for 1 min). Homogenates were clarified by centrifugation (16800 × g for 15 min at 4°C) and stored at −80 °C until analysis.

### 2.5 Tissue infectious virus quantification

E5 or CVB3 infectious virus titers in whole blood or tissue homogenate were determined by TCID_50_ assay^26; 38^. Samples were serially diluted in HeLa media and plated onto a monolayer of confluent HeLa cells in replicates of 6. Cells were incubated for 5 days at 37°C and 5% CO2, fixed > 1 h with 4% formaldehyde, and then stained with crystal violet for 20 min. After staining, plates were scored for cytopathic effects and the Reed-Muench method^39^ was used to calculate the TCID_50_ titer for each sample.

### 2.6 Anti-E5 ELISA

Anti-E5 antibody binding titers in serum were measured using an established protocol^38; 40; 41^, with modifications. Microlon 96 well high binding plates (Greiner Bio-One #655081) were coated with 100ng per well of purified inactivated E5 (described above) in pH 9.6 carbonate buffer overnight at 4°C. Plates were washed with wash buffer [PBS plus 0.1% Tween-20 (Sigma #P1379)] and blocked for 1 h at 37°C with 10% nonfat milk (RPI #M17200) in PBS. After washing, samples were serially diluted (starting at 1:100) in wash buffer plus 10% nonfat milk and 10% bovine serum albumin (BSA; RPI #A30075-100) and added in duplicate for 1 h at 37°C. Following washing, horseradish peroxidase (HRP)-conjugated secondary antibodies, specific for the isotype of interest, were diluted in wash buffer and added for 1 h at 37°C. The following secondary antibodies were used: anti-mouse total IgG (predicted to detect mouse and chimeric IgG; Invitrogen #32430, 1:250), anti-human IgG Fcγ (to detect chimeric IgG1^42^; Jackson ImmunoResearch #109-035-008, 1:5000), anti-mouse IgG2c (Cell Signaling #56970, 1:20000), anti-mouse IgM (Invitrogen # 62-6820, 1:2000), and anti-mouse IgA (Invitrogen #62-6720, 1:2000) The plates were then washed, developed with 3,3’,5,5’ tetramethylbenzidine (TMB; BD Biosciences # 55214) for 15-20 min, and stopped using 1N hydrochloric acid. Plates were read at 450 nm absorbance on a SpectraMax iD3 plate reader (Molecular Devices). Antibody titers were defined as the highest serum dilution with an OD value above the cutoff, calculated as the mean of negative controls plus 3 standard deviations per experiment. All washes were performed with a BioTek 405 TS plate washer (Agilent).

### 2.7 Microneutralization assay

Anti-E5 neutralizing antibody titers were measured in serum by tissue culture microneutralization assays^38; 40; 41^. Serum samples were heat inactivated at 56°C for 30 min on a heat block. Samples were serially diluted in HeLa media starting at 1:20 then mixed with 100 TCID_50_ units of E5 and incubated for 1 h at room temperature. The mixture was used to infect duplicate wells of confluent HeLa cells for 1 hour at 37°C. After incubation, the inoculum was removed, cells were washed once with PBS, and fresh HeLa media was added. Cells were incubated for 3 days at 37°C and 5% CO2, fixed for >1 h with 4% formaldehyde, and stained with crystal violet for 20 min. Neutralizing antibody titers were calculated as the highest serum dilution that prevented virus cytopathic effects in 50% of wells.

### 2.8 Passive serum transfer

Male and female hFc mice (6-10 weeks of age) were infected IP with 10^3^ PFU of E5 or mock-inoculated, monitored, and bled for serum samples at day 42 post-infection as described above. Heat-inactivated (56°C for 30 min) serum from E5-infected and mock-inoculated mice was pooled by infection status, and the presence or absence, respectively, of anti-E5 neutralizing antibodies was confirmed prior to passive transfer. Naïve male and female hFc mice (8-10 weeks of age) were injected IP with 200 μL of pooled heat-inactivated serum from either E5-infected or mock-inoculated mice. Twenty-four hours later, mice were challenged IP with 10^6^ PFU of E5 and monitored for 14 days as described above.

### 2.9 Statistical analysis

The area under the curve (AUC) of body mass change and log2-transformed neutralizing antibody titer curves, log10-transformed virus titers, and log2-transformed antibody titers were analyzed with two-way analysis of variance (ANOVA) followed by *post-hoc* Tukey multiple comparisons tests. Kaplan-Meier survival curves were analyzed with the Mantel-Cox Logrank test. Associations between specific antibody titers and survival were assessed using Firth’s penalized likelihood logistic regression models^43^. Log2-transformed titers were used as the primary predictor, and biological sex was included as a modifying variable. This approach allows the relationship between antibody titers and survival to differ by sex, enabling estimation of sex-specific models, rather than assuming a single effect for both sexes.^44^ Model coefficients, 95% confidence intervals, and coefficient p values are reported. Model performance was evaluated by receiver operating characteristic (ROC) analysis, with AUC as the measure of model predictive accuracy. Predictive titers associated with >90% probability of survival were derived from fitted models with statistically significant coefficients for male and female mice, respectively. For ease of interpretation, predictive titers (log2) were converted back to reciprocal dilutions (e.g., 1:50, 1:100) corresponding to the nearest 2-fold assay dilution when reported in the text. Statistical significance was defined as p < 0.05. Analyses were performed using GraphPad Prism (v10.6) and R (v4.5.1) using the logistf and pROC packages. R code used is available at CoyneLabDuke GitHub repository at https://github.com/CoyneLabDuke/Echovirus-Antibodies.

## 3 Results

### 3.1 E5 infection of hFc mice results in acute disease and elicits robust virus-specific neutralizing antibodies

Tg32-hFcRn (Tg32) mice, which lack mouse FcRn and express human FcRn (the pan-echovirus receptor^23^) under the native promoter, are an established model of acute echovirus pathogenesis^26; 27; 34^, but exhibit impaired antibody responses due to an inability to recycle mouse IgG^32^. To address this limitation, Tg32 mice have been further engineered to produce chimeric IgG1 with mouse Fab regions and the human Fc domain.^32^ We hypothesized that these modified mice (hereafter referred to as hFc mice) would have improved antibody responses following echovirus 5 (E5) infection. Because biological sex can influence viral pathogenesis and immune responses^45–47^, including in enterovirus infections^48–50^, both male and female mice were included and evaluated independently. Following sublethal-dose intraperitoneal (IP) E5 infection, which models disseminated infection^27; 34^, both male and female hFc mice experienced morbidity (measured by body mass loss; Fig. 1A-B) and survival outcomes (Fig. 1C) equivalent to Tg32 controls. Infectious E5 titers at 3 days post-infection (dpi) in blood (Fig. 1D) as well as the liver (Fig. 1E) and pancreas (Fig. 1F), two key sites of acute echovirus replication^27^, did not differ between hFc and Tg32 mice, consistent with acute echovirus replication being controlled by innate rather than antibody-mediated responses^27; 34^.

**Figure 1.**
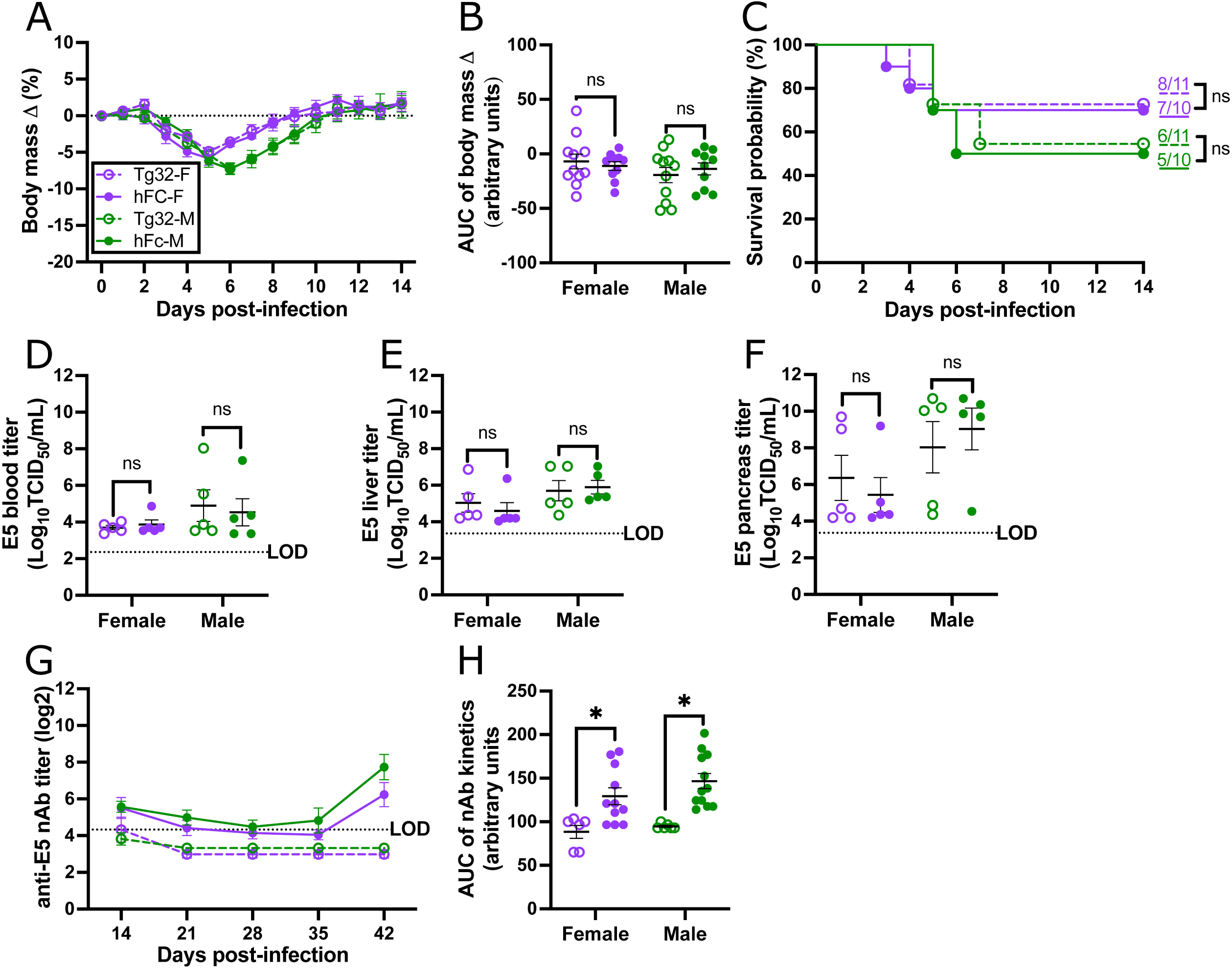
Echovirus 5 (E5) infection of hFc mice results in acute infection and elicits robust virus-specific neutralizing antibodies (nAb). Male and female Tg32 (open circles) and hFc (closed circles) mice were intraperitoneally infected with 10^3^ PFU of E5 or mock-inoculated with PBS and monitored for changes in body mass (A), with the area under the curve (AUC) recorded as a measure of the difference in mass change over time (B). Survival after E5 infection was documented and compared via Kaplan-Meier plot (C). A subset of infected mice was euthanized at 3 days post-infection, with blood (D), liver (E), and pancreas (F) collected and infectious virus measured by TCID_50_ assay. Surviving mice were bled at 14, 21, 28, 35, and 42 days-post infection, serum collected, and anti-E5 nAb titers measured by microneutralization assay (G), with AUC recorded as a measure of the nAb response over time (H). Circles (A,G), or bars (B, D, E, F, H) represent the mean ± standard error of the mean from two independent replications [n = 10–11 (A-C), 5 (D-F), or 6-12 (G-H) mice/group] with individual mice indicated by circles (B, D, E, F, H). Significant differences (p < 0.05) were determined by two-way ANOVA with Tukey post hoc test or Mantel-Cox logrank. The limit of detection (LOD) is indicated with a dashed line. Asterisk (*) indicates statistically significant differences and ns indicates not significant.

We next assessed whether hFc mice had improved antibody responses following infection by serially sampling Tg32 and hFc mice weekly and measuring E5-specific nAb titers (Fig. 1G-H). Tg32 mice had limited nAb titers at 14 dpi, which fell below the limit of detection at later timepoints, consistent with an inability of Tg32 mice to recycle mouse IgG with human FcRn^32^. In contrast, hFc mice had detectable nAb titers at 14 dpi that further increased at 42 dpi (Fig. 1G), with a significantly greater nAb response overall (as measured by AUC of the nAb titer kinetics) compared to Tg32 mice (Fig. 1H). Taken together, these data demonstrate that hFc mice mount a functional antibody response following E5 infection without altering acute pathogenesis, providing a mouse model to characterize the isotype distribution and protective capacity of E5-specific antibodies.

### 3.2 E5-infected hFc mice produce durable neutralizing antibodies as well as virus-specific IgG and IgM

Because hFc mice had superior nAb production compared to the parental Tg32 line (Fig. 1G-H), we used hFc mice for all subsequent experiments. Male and female hFc mice were mock- or E5-infected IP, and serum collected at 14 and 42 dpi for antibody analysis (Fig. 2A). To characterize the breadth of the antibody response, we measured E5-specific nAb titers (Fig. 2B, H) and E5-specific titers of total IgG (Fig 2C, I), chimeric IgG1 (Fig. 2D, J), mouse IgG2c (the predominant IgG subtype in C57BL/6-background mice following infection^51–53^; Fig. 2E, K), mouse IgM (Fig. 2F, L), and mouse IgA (Fig. 2G, M).

**Figure 2.**
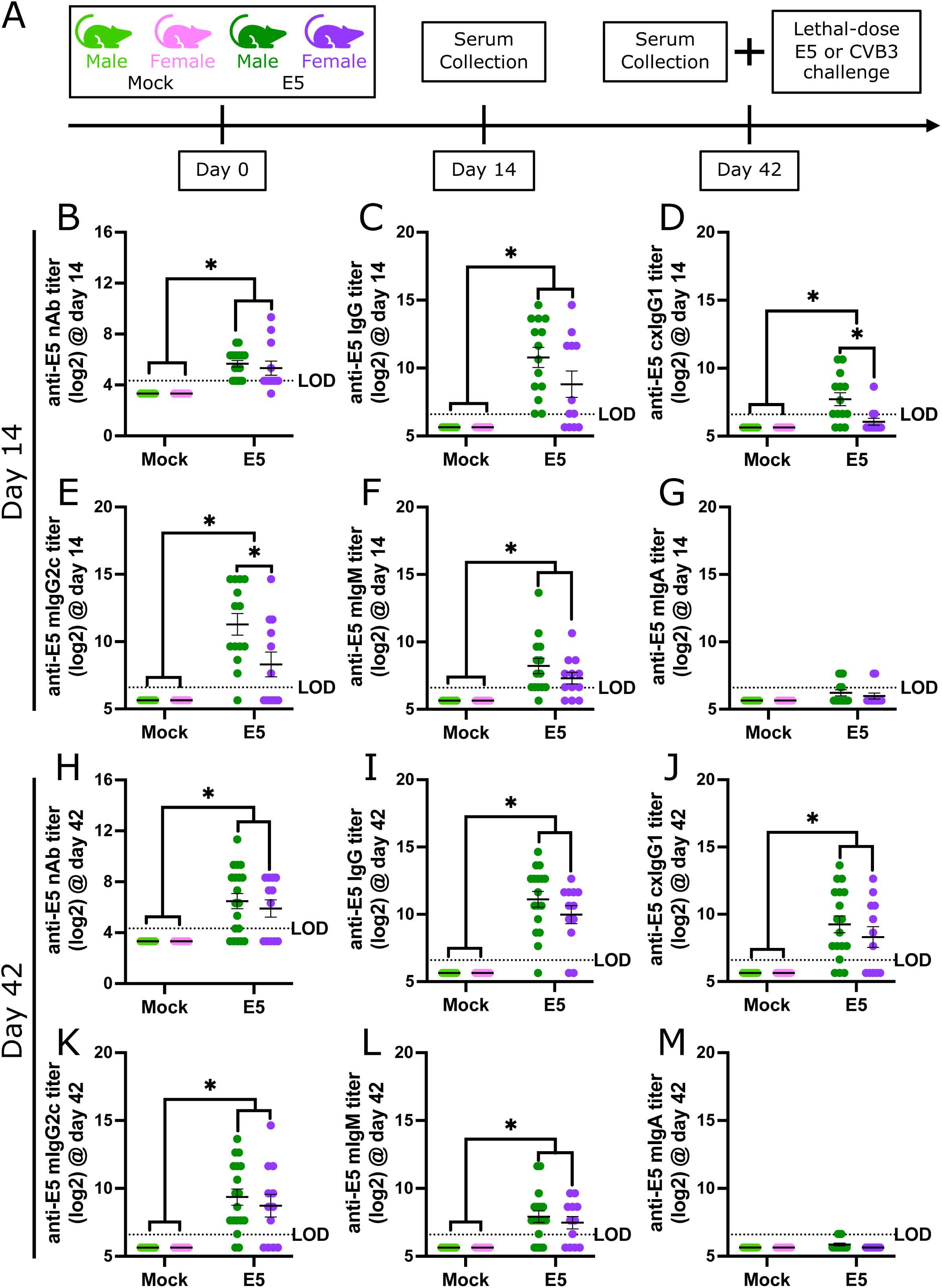
Echovirus 5 (E5)-infected hFc mice produce durable neutralizing antibodies (nAb) as well as virus-specific IgG and IgM. Male and female hFc mice were intraperitoneally infected with 10^3^ PFU of E5 or mock-inoculated with PBS, bled for serum collection at day 14 and 42, and challenged with 10^6^ PFU of E5 or CVB3 (A). In day 14 (B-G) and 42 (H-M) serum, anti-E5 nAb titers were measured by microneutralization assay (B, H), and anti-E5 total IgG (C, I), chimeric (cx) IgG1 (D, J), mouse (m) IgG2c (E, K), mIgM (F, L), and mIgA (G, M) were measured by ELISA. Bars represent the mean ± standard error of the mean from two independent replications (n = 8–19mice/group) with individual mice indicated with circles. Antibody titers were log2-transformed for analysis. Significant differences (p < 0.05) were determined by two-way ANOVA with Tukey post hoc test. The limit of detection (LOD) is indicated with a dashed line. Asterisk (*) indicates statistically significant differences.

At 14 dpi, E5 infected mice developed appreciable anti-E5 nAb titers (Fig. 2B) as well as anti-E5 total IgG (Fig. 2C), IgG1 (Fig. 2D), IgG2c (Fig 2E), and IgM (Fig. 2F) titers. At 42 dpi, antibody responses were robust, with high mean titers of nAb (Fig. 2H), total IgG (Fig. 2I), chimeric IgG1 (Fig. 2J), mouse IgG2c (Fig. 2K) and IgM (Fig. 2L) in previously infected mice. In contrast, serum anti-E5 IgA titers were low or undetectable at both time points (Fig. 2G, M). Transient sex differences were observed, with E5-infected males having more IgG1 and IgG2c than females at 14 dpi (Fig. 2D, E) but not 42 dpi (Fig. 2J, K). These data indicate that E5 infection of hFc mice results in a robust antibody response in both males and females.

### 3.3 Prior E5 infection confers partial, type-specific protection against lethal challenge

Given the induction of E5-specific antibody responses, we next asked whether prior infection confers protection against reinfection. To test this, we challenged mock- or E5-infected hFc mice at 42 dpi with a lethal dose of E5. Previously mock-inoculated male and female mice experienced significant morbidity, measured by loss in body mass (Fig. 3A-B), and all succumbed to infection within 7 days-post challenge (dpc; Fig. 3C). In contrast, mice previously infected with E5 were partially protected, with reduced morbidity (Fig. 3A-B) as well as survival in approximately 75% of males and 55% of females (Fig. 3C). To assess whether protection included sterilizing immunity, a subset of mice was collected at 3 dpc to measure viral replication in the liver, a major site of echovirus replication^27^ (Fig 3D). Compared to previously-mock-inoculated mice, E5-infected male and female mice had significantly reduced liver titers. In some mice, infectious virus was below the limit of detection, consistent with sterilizing immunity.

**Figure 3.**
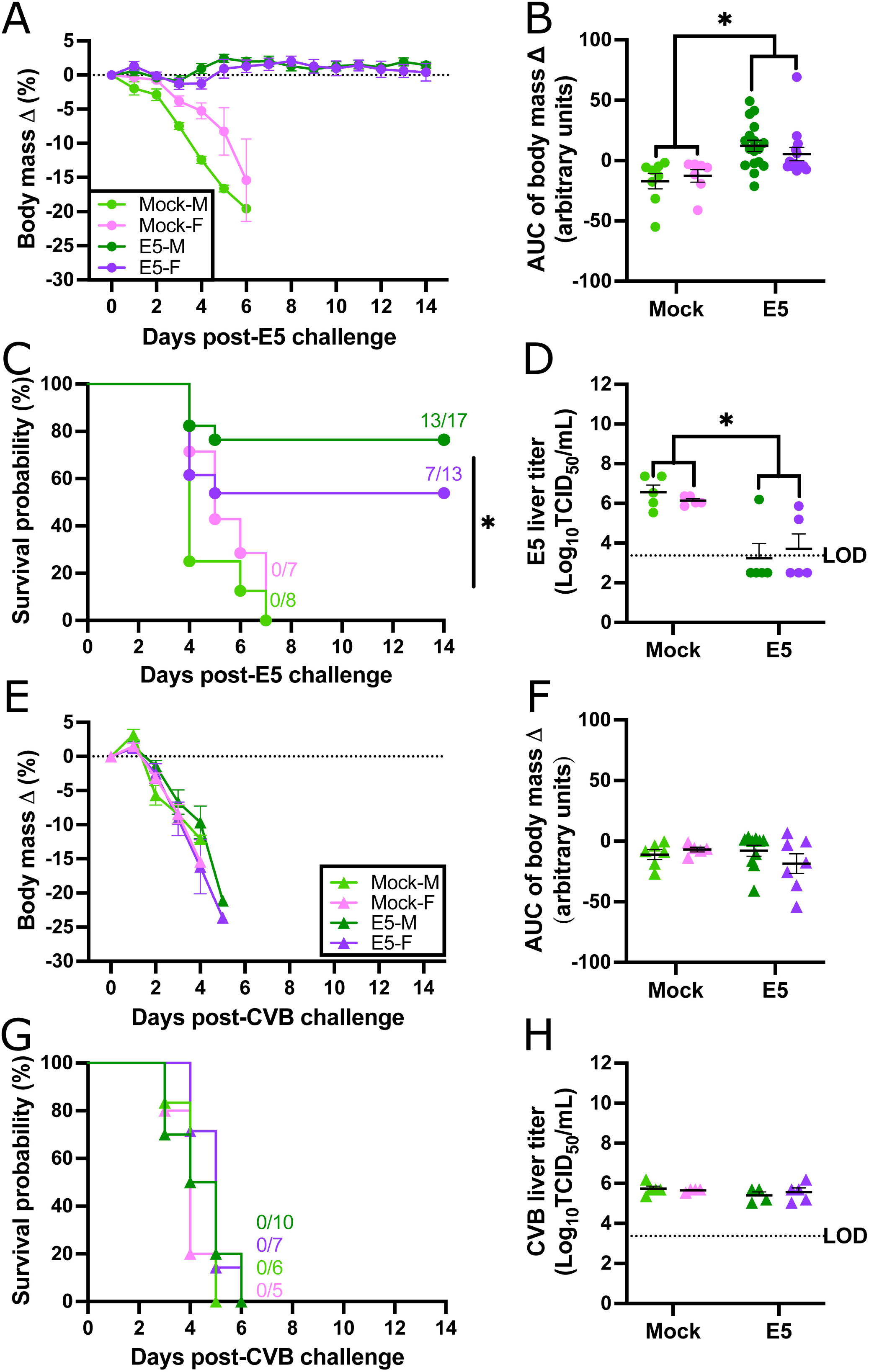
Prior E5 infection confers partial, type-specific protection against lethal challenge. Male and female hFc mice that were previously infected with 10^3^ PFU of E5 or mock-inoculated with PBS were challenged with 10^6^ PFU of E5 (A-D) or CVB3 (E-H) and monitored for changes in body mass (A, E), with the area under the curve (AUC) recorded as a measure of the difference in mass change over time (B, F). Survival after challenge was documented and compared via Kaplan-Meier plot (C, G). A subset of challenged mice were euthanized at 3 days post-challenge, liver collected, and infectious virus measured by TCID_50_ assay (D, H). Circles (A,E) or bars (B, D, F, H) represent the mean ± standard error of the mean from two independent replications [n = 7-13 (A-C), 5 (D, H), or 5-10 (E-G) mice/group] with individual mice indicated by circles (B, D, F, H). Significant differences (p < 0.05) were determined by two-way ANOVA with Tukey post hoc test or Mantel-Cox logrank. The limit of detection (LOD) is indicated with a dashed line. Asterisk (*) indicates statistically significant differences.

Because enterovirus immunity is generally considered to be type-(traditionally referred to as serotype^54^) specific, with antibodies showing little to no cross-type neutralization^19; 55–57^, we next tested protection against a heterotypic challenge. Mock- and E5-infected mice were infected at 42 dpi with CVB3, a closely related enterovirus B. Previous E5 infection had no significant effect on CVB3 morbidity (Fig. 3E-F) or mortality (Fig. 3G). Liver titers at 3 dpc were similarly high in all groups (Fig. 3H), confirming the absence of cross-protection. Taken together, these data indicate that sublethal echovirus infection results in type-specific protection, with survival and sterilizing immunity in a subset (>50%) of mice but not extending to heterotypic enteroviruses.

### 3.4 Serum neutralizing antibodies at the time of challenge correlate with survival

Because protection from E5 reinfection was partial, we next evaluated whether antibody titers could predict survival following challenge. To do this, we reorganized antibody titers of E5-infected mice according to whether mice ultimately survived (+) or succumbed (-) to E5 challenge (Fig 4). At day 14, there were no significant differences between survivors and non-survivors in anti-E5 nAb titers (Fig. 4A) or any isotype measured (Fig. 4B-F). At day 42, the day of challenge, surviving male and female mice had significantly higher anti-E5 nAb (Fig. 4G), total IgG (Fig. 4H), and IgG1 (Fig. 4I) titers, whereas only females had significantly higher IgG2c (Fig. 4J), and IgM (Fig. 4K) titers compared to non-survivors. These data suggest that serum neutralizing antibodies correlate with survival regardless of sex, but the contributing isotypes may differ between males and females.

**Figure 4.**
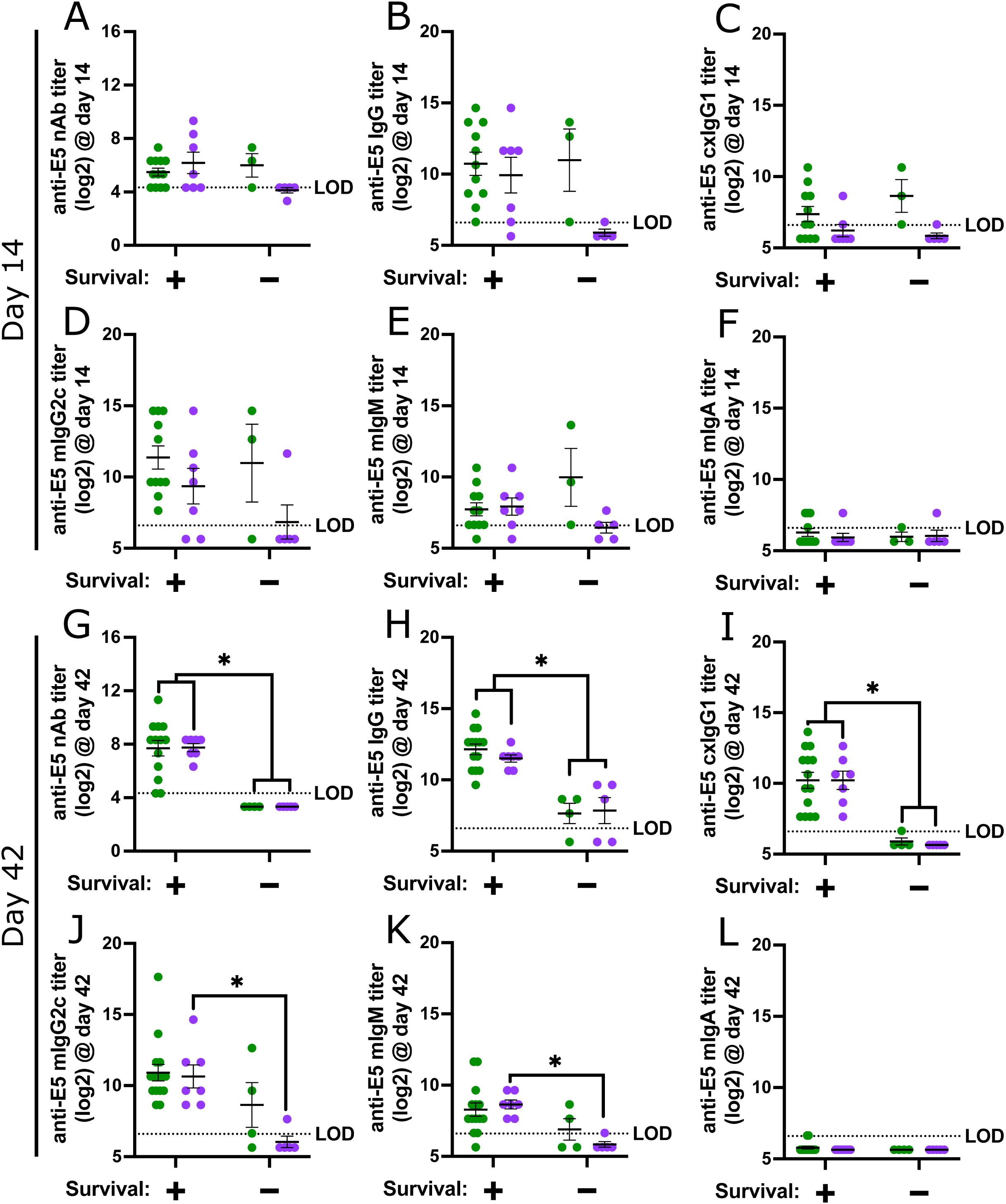
Serum anti-echovirus 5 (E5) neutralizing antibody (nAb), IgG, and IgM titers at the time of challenge distinguish survivors of E5 challenge. Serum log2-transformed anti-E5 nAb (A, G), total IgG (B, H), chimeric (cx) IgG1 (C, I), mouse (m) IgG2c (D, J), mIgM (E, K), and mIgA (F, L) titers at 14 (A-F) and 42 (G-L) days post-E5 infection from Figure 2 were reorganized according to whether mice ultimately survived (+) or succumbed (-) to E5 challenge. Bars represent the mean ± standard error of the mean from two independent replications (n = 3-14 mice/group) with individual mice indicated with circles. Significant differences (p < 0.05) were determined by two-way ANOVA with Tukey post hoc test. The limit of detection (LOD) is indicated with a dashed line. Asterisk (*) indicates statistically significant differences.

To quantitatively identify antibody titers associated with survival^58^, we used Firth’s logistic regression models (Table 1), which are preferred for preclinical studies with small datasets^43^. Log2-transformed antibody titers at 14 (Table 1A) or 42 dpi (Table 1B) were included as the primary predictor in each model, with sex included as a modifying variable. Although sex was not an independent predictor of survival, this approach allowed estimation of predictive titers for both males and females^44^. At 42 dpi, anti-E5 nAb, total IgG, and chimeric IgG1 titers were most predictive of survival, as indicated by receiver operating characteristic (ROC) AUC values of 1 for each model. In males, the models predict >90% probability of survival with log2 nAb titers ≥ 5.9 (∼1:40), total IgG titers ≥ 10.7 (∼1:1600), and chimeric IgG1 titers ≥ 7.8 (∼1:200). In females, the corresponding predictive log2 titers were ≥ 6.5 (∼1:80) for nAb, ≥ 11.2 (∼1:1600) for total IgG, and ≥ 7.9 (∼1:200) for chimeric IgG1. Anti-E5 IgG2c and IgM at 42 dpi were also significant predictors, but with lower predictive ability than nAb, total IgG, or chimeric IgG1, as reflected by lower ROC AUC values. In contrast, no antibody measures at 14 dpi were predictive of survival. Taken together, these results indicate that serum neutralizing antibodies, total IgG, and IgG1 at the time of challenge serve as quantitative correlates of protection from lethal E5 infection and highlight potential immunological targets for future echovirus vaccine development^58^.

**Table 1.**
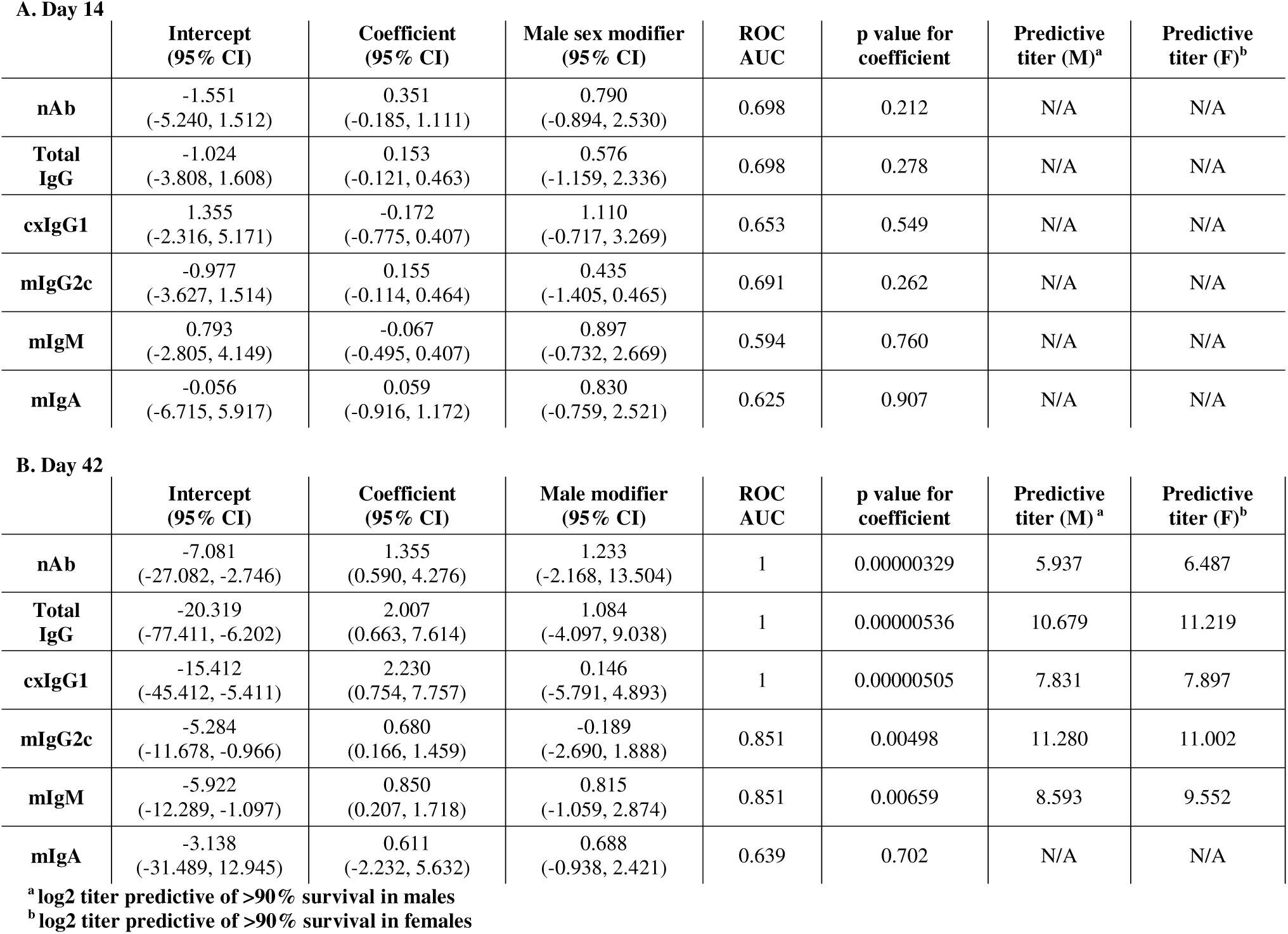
Firth’s logistic regression models of antibody titers at day 14 and day 42 post infection.

| A. Day 14 |  |  |  |  |  |  |  |
| --- | --- | --- | --- | --- | --- | --- | --- |
|  | Intercept<br>(95% CI) | Coefficient<br>(95% CI) | Male sex modifier<br>(95% CI) | ROC<br>AUC | p value for<br>coefficient | Predictive<br>titer (M)<sup>a</sup> | Predictive<br>titer (F)<sup>b</sup> |
| nAb | -1.551<br>(-5.240, 1.512) | 0.351<br>(-0.185, 1.111) | 0.790<br>(-0.894, 2.530) | 0.698 | 0.212 | N/A | N/A |
| Total<br>IgG | -1.024<br>(-3.808, 1.608) | 0.153<br>(-0.121, 0.463) | 0.576<br>(-1.159, 2.336) | 0.698 | 0.278 | N/A | N/A |
| cxIgG1 | 1.355<br>(-2.316, 5.171) | -0.172<br>(-0.775, 0.407) | 1.110<br>(-0.717, 3.269) | 0.653 | 0.549 | N/A | N/A |
| mIgG2c | -0.977<br>(-3.627, 1.514) | 0.155<br>(-0.114, 0.464) | 0.435<br>(-1.405, 0.465) | 0.691 | 0.262 | N/A | N/A |
| mIgM | 0.793<br>(-2.805, 4.149) | -0.067<br>(-0.495, 0.407) | 0.897<br>(-0.732, 2.669) | 0.594 | 0.760 | N/A | N/A |
| mIgA | -0.056<br>(-6.715, 5.917) | 0.059<br>(-0.916, 1.172) | 0.830<br>(-0.759, 2.521) | 0.625 | 0.907 | N/A | N/A |
| B. Day 42 |  |  |  |  |  |  |  |
|  | Intercept<br>(95% CI) | Coefficient<br>(95% CI) | Male modifier<br>(95% CI) | ROC<br>AUC | p value for<br>coefficient | Predictive<br>titer (M)<sup>a</sup> | Predictive<br>titer (F)<sup>b</sup> |
| nAb | -7.081<br>(-27.082, -2.746) | 1.355<br>(0.590, 4.276) | 1.233<br>(-2.168, 13.504) | 1 | 0.00000329 | 5.937 | 6.487 |
| Total<br>IgG | -20.319<br>(-77.411, -6.202) | 2.007<br>(0.663, 7.614) | 1.084<br>(-4.097, 9.038) | 1 | 0.00000536 | 10.679 | 11.219 |
| cxIgG1 | -15.412<br>(-45.412, -5.411) | 2.230<br>(0.754, 7.757) | 0.146<br>(-5.791, 4.893) | 1 | 0.00000505 | 7.831 | 7.897 |
| mIgG2c | -5.284<br>(-11.678, -0.966) | 0.680<br>(0.166, 1.459) | -0.189<br>(-2.690, 1.888) | 0.851 | 0.00498 | 11.280 | 11.002 |
| mIgM | -5.922<br>(-12.289, -1.097) | 0.850<br>(0.207, 1.718) | 0.815<br>(-1.059, 2.874) | 0.851 | 0.00659 | 8.593 | 9.552 |
| mIgA | -3.138<br>(-31.489, 12.945) | 0.611<br>(-2.232, 5.632) | 0.688<br>(-0.938, 2.421) | 0.639 | 0.702 | N/A | N/A |

<sup>a</sup> log<sub>2</sub> titer predictive of >90% survival in males<sup>b</sup> log<sub>2</sub> titer predictive of >90% survival in females

### 3.5 Passive transfer of serum with anti-E5 nAbs protects naïve mice from E5 challenge

Having identified serum anti-E5 nAbs as a correlate of protection, we next tested whether these antibodies are sufficient to confer protection. To do so, naïve mice received pooled heat-inactivated serum from either E5-immune or mock-inoculated mice and were subsequently challenged with a lethal dose of E5 (Fig. 5A). Both male and female mice injected with pooled E5-immune serum were completely protected from morbidity (Fig. 5B-C) and mortality (Fig. 5D), in contrast to mock-serum controls, which succumbed to infection by 7 dpc. Furthermore, E5-immune serum prevented virus replication in the liver, as titers were undetectable in recipients of E5-immune serum, whereas high virus titers were observed in mock-serum controls (Fig. 5E). These data demonstrate that serum containing anti-E5 antibodies are sufficient to protect naïve mice from lethal challenge and further support anti-E5 nAbs as a mechanistically relevant correlate of protection.

**Figure 5.**
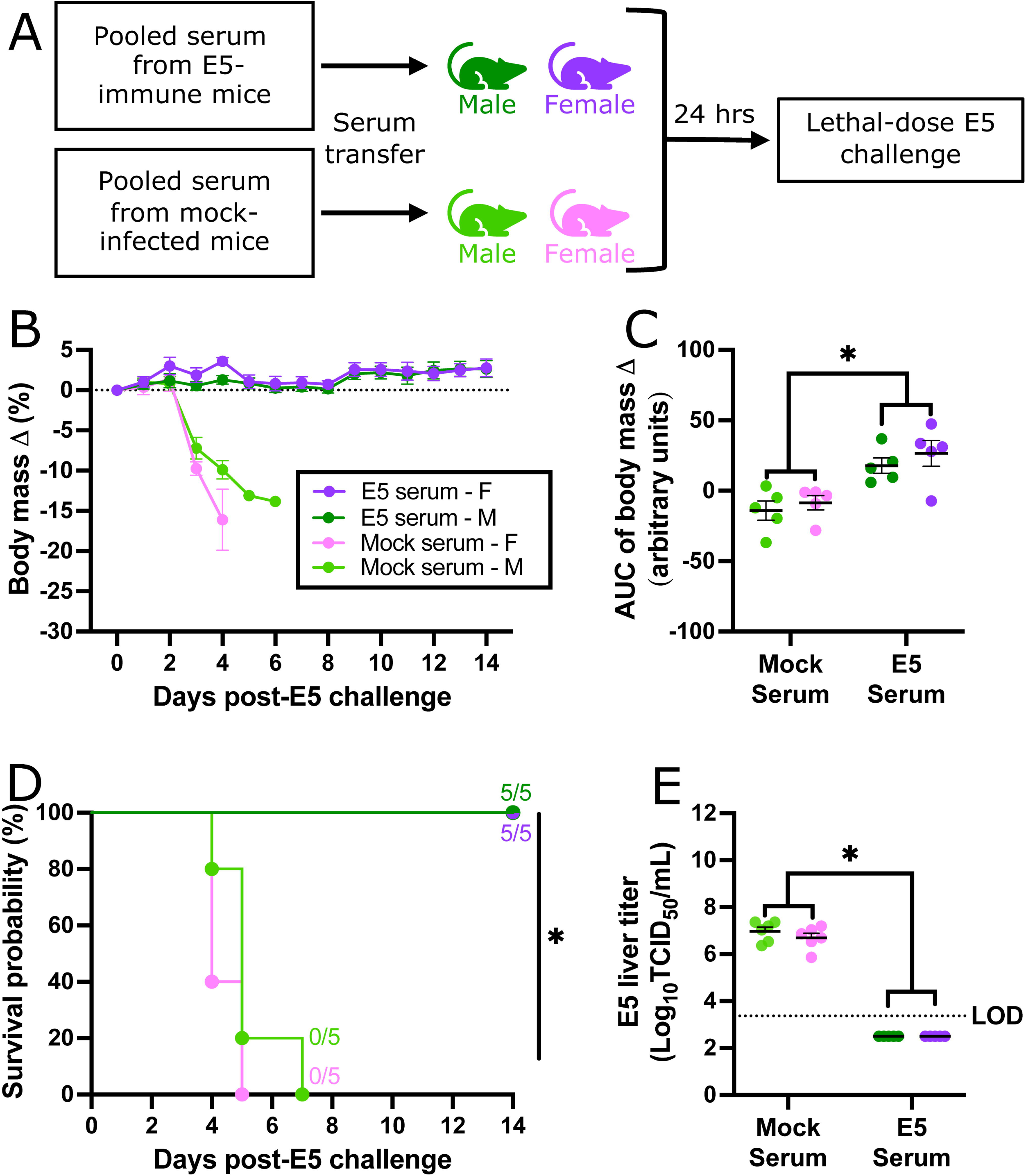
Passive transfer of serum with anti-echovirus 5 (E5) neutralizing antibodies (nAb) protects naïve mice from E5 challenge. Heat-inactivated serum from E5-infected or mock-inoculated mice was pooled by immune status and injected intraperitoneally into naïve adult male and female hFc mice. Twenty-four hours later, mice were challenged IP with 10^6^ PFU of E5 (A) and monitored for changes in body mass (B), with the area under the curve (AUC) recorded as a measure of the difference in mass change over time (C). Survival after challenge was documented and compared via Kaplan-Meier plot (D). A subset of challenged mice were euthanized at 3 days post-challenge, liver collected, and infectious virus measured by TCID_50_ assay (E). Circles (B), or bars (C,E) represent the mean ± standard error of the mean from two independent replications (n = 5 mice/group) with individual mice indicated by circles (C, E). Significant differences (p < 0.05) were determined by two-way ANOVA with Tukey post hoc test or Mantel-Cox logrank. The limit of detection (LOD) is indicated with a dashed line. Asterisk (*) indicates statistically significant differences.

## 4 Discussion

The absence of a small animal model has traditionally limited research into *in vivo* immune responses to echoviruses. The discovery of human FcRn as a pan-echovirus receptor and the subsequent use of Tg32 human FcRn mice advanced the field by enabling the study of innate immune mechanisms and echovirus pathogenesis *in vivo*^23; 27^. Building on this foundation, the current study utilized hFc mice, which are Tg32 mice further engineered to produce chimeric IgG1 that can be recycled by human FcRn, restoring circulating IgG^32^. We found that echovirus-infected hFc mice exhibited acute pathogenesis comparable to the parental Tg32 line, while also mounting a more robust and durable humoral response, validating their use as a translational platform for immunological studies of echovirus infection.

Primary echovirus infection of hFc mice elicited appreciable virus-specific serum nAbs early in recovery and at the time of challenge. Notably, while most mice developed detectable titers, a subset did not, and this heterogeneity was reflected in survival outcomes following challenge. We leveraged this natural variation to show that the presence of echovirus-specific nAbs, total IgG, and chimeric IgG1 were most strongly associated with protection. Moreover, passive transfer of immune serum from previously infected mice to naïve recipients conferred complete, sterilizing protection against morbidity and mortality. Together, these findings demonstrate that virus-specific nAbs are the dominant driver of protection from echovirus reinfection *in vivo*, consistent with clinical observations in humans, where patients with impaired antibody responses (i.e., genetic agammaglobulinemia or B cell-depletive therapy) are highly susceptible to severe echovirus infection^29; 30^. Placed in a broader context, our results suggest that adaptive immunity to echoviruses may mirror that of other enteroviruses such as CVB3 and EV-A71, where antibody production is a near absolute requirement for protection.^59; 60^

Enterovirus serology has traditionally relied on nAbs to define exposure patterns, as nAbs induced by enterovirus infections are generally type-specific, *in vitro*^61; 62^. However, some enterovirus antibodies can exhibit non-neutralizing, cross-type binding^55; 63; 64^, raising the question of whether antibody-mediated protection following echovirus infection is strictly type-specific *in vivo*. In the current study, prior E5 infection failed to protect hFc mice against challenge with the genetically closely related, heterotypic CVB3, demonstrating limited cross-type protection, consistent with observations from monovalent EV-A71 vaccines which also confer type-specific protection^65^. Nevertheless, E5 nAbs may be cross-protective against other enteroviruses not tested here, as some enterovirus-specific antibodies have limited cross-type neutralization against genetically distant enteroviruses^55^. In contrast to other viral infections where CD8+ T cells play a key protective role^66^, enteroviruses generally induce weak cytotoxic T cell responses, which contribute minimally to protection^67; 68^. Although T cell responses were not measured in this study, the lack of heterotypic protection suggests a limited contribution of cytotoxic T cells, as these cells generally target conserved antigens and would be expected to provide partial cross-protection^69^. Future studies, such as generating B- or T-cell deficient hFc mice will be critical to dissect the cellular contributions to echovirus adaptive immunity.

Effective therapeutics for echovirus infection are limited. Intravenous immunoglobulin (IVIG) has traditionally been used, particularly in neonates and immunocompromised individuals, but its efficacy is inconsistent ^70; 71^, likely due to lot-to-lot variability in the concentration of nAbs specific to the patient’s infecting echovirus. For example, IVIG with a neutralization titer of ≥1:800 to a patient’s viral isolates has been associated with more rapid cessation of viremia, whereas IVIG with lower neutralization titers was less effective^71^. Our findings support this mechanism, as virus-specific nAbs were strongly associated with protection *in vivo*, potentially explaining why IVIG lacking relevant nAbs may be ineffective. Moreover, new therapeutics with consistent efficacy against echoviruses are needed, and the hFc model of echovirus infection provides a valuable preclinical platform to test these promising therapies *in vivo*.

Correlates of protection are critical for guiding vaccine design and development^58; 72^. For example, hemagglutination inhibition (HAI) antibody titers of approximately ≥1:40 are an established a correlate of protection for influenza vaccines, associated with a roughly 50% reduction in risk for symptomatic influenza^73; 74^. Defined correlates improve vaccine studies by providing a measurable and standardized immunological endpoint, informing dosing decisions, and enabling cross-platform comparisons^58; 75^. In this study, we used Firth’s penalized logistic regressions to identify antibody titers associated with protection. We calculated these thresholds based on a survival rate of > 90%, but they can be adjusted to any desired protective level, providing a flexible framework for defining protective antibody titer thresholds in future echovirus vaccine studies. Small animal models of infection, such as hFc mice, enable preclinical evaluation of vaccine efficacy with lethal-dose challenge. Together, our work establishes a translational platform that links protective humoral responses to defined correlates of protection, supporting the development of echovirus vaccines and therapeutics while improving our mechanistic understanding of adaptive immunity to echoviruses.

## Acknowledgements

The authors would like to thank Dr. J Lindsay Whitton (The Scripps Research Institute) for providing the CVB3 pH3 infectious clone and the expert animal care staff of Duke University’s Division of Laboratory Animal Resources for assistance with maintaining our mouse colony.

## 6 Funding

This project was supported by NIH/NIAID 1U19AI181979-01 (C.B.C)

## 7 Conflicts of Interest

None declared.

## 8 Author contributions

Conceptualization: P.S.C., C.B.C.; Data curation: P.S.C., J.W.A.; Formal analysis: P.S.C., J.W.A.; Funding acquisition: C.B.C.; Investigation: P.S.C., J.W.A.; Methodology: P.S.C., C.B.C.; Project administration: P.S.C., C.B.C.; Resources: P.S.C., K.A.C., C.B.C.; Supervision: C.B.C.; Validation: P.S.C., J.W.A.; Visualization: P.S.C.; Writing – original draft: P.S.C., C.B.C.; Writing – review & editing: P.S.C., C.B.C., J.W.A., K.A.C.

## 9 Data availability

R code used and the associated data is publicly available at CoyneLabDuke GitHub repository at https://github.com/CoyneLabDuke/Echovirus-Antibodies.

## Notes

### Competing Interest Statement

The authors have declared no competing interest.

